# A spinal origin for the obligate flexor synergy in the non-human primate: Implications for control of reaching

**DOI:** 10.1101/2025.07.28.666086

**Authors:** Isabel S. Glover, Anne M.E. Baker, John W. Krakauer, Stuart N. Baker

## Abstract

Stroke survivors frequently develop the flexor synergy, an obligate co-contraction of shoulder abductors and elbow flexors; the neural substrate has proven elusive. Here we trained healthy monkeys to generate isometric elbow and shoulder torques to move an on-screen cursor, and recorded neuron firing from motor cortical areas and the reticular formation. In all regions we found cells correlated with activity around a single joint. Neurons coding co-contractions showed a bias towards combinations orthogonal to the post-stroke flexor synergy, e.g. shoulder abduction with elbow extension. Threshold microstimulation in the spinal cord but not in either motor cortex or the reticular formation generated coactivation aligned to the flexor synergy. We suggest the evolution of prehension required descending systems either to control or bypass locomotion-dedicated spinal circuits. Loss of descending input after stroke constrains the upper limb to spinal synergies best suited to primitive locomotor functions.

## Introduction

After a stroke involving motor cortex and its corticofugal outputs, patients typically pass through a defined sequence towards recovery^1, 2^. There is an initial flaccid paralysis, caused by the loss of descending input to motoneurons. This is then replaced by weak and awkward movements characterized by synergies – obligate co-contractions of muscles acting around different joints. The most common is the flexor synergy, in which abduction around the shoulder joint is coupled with flexion around the elbow, wrist and digits^3-5^. With further recovery, a patient may be able to escape the constraint of synergies and make more normal movements. However, this is not always possible; for the most severely affected stroke survivors, synergies remain a permanent feature of their movements and a major contributor to disability^6, 7^. They are therefore an important target for rehabilitation.

Why should damage to the motor cortex lead to synergies? One plausible idea is that they are a release phenomenon, produced by a motor circuit which naturally generates the observed co-activation and which is normally suppressed in healthy individuals. For example, in the newborn, Twitchell ^8^ described the traction response which has a similar pattern of muscle activation to the flexor synergy; it is usually lost over the first four months of life. This dormant circuit could reemerge post-stroke and cause synergies. It is also possible that the underlying circuit plays an everyday role in movements of the healthy adult, being activated for movements which require muscle patterns coincident with the synergy. In health, this circuit would be activated or inhibited by cortical systems as appropriate to the task at hand. However, the loss of cortical control after a stroke could lead to a failure of inhibitory control, and the inappropriate intrusion of synergy coactivations into movements where they are unwanted. For example, activation of the flexor synergy during reaching away from the body leads to a severe limitation of available workspace^9^, because coactivation of elbow flexion with shoulder abduction is the opposite of the elbow extension and shoulder abduction required.

Where in the central nervous system is the circuit for synergies? The available evidence suggests a role for the contralesional motor cortex, and its indirect connections to the paretic limb via the reticular formation in the brainstem and the reticulospinal tract^10, 11^. It is known that the reticulospinal tract is less able to activate muscles selectively than the corticospinal tract^12, 13^. Neurons in the reticular formation become more active after a corticospinal lesion^14^, and reticulospinal connections strengthen^15^. Increased reliance on this pathway is therefore a plausible route to produce the obligate muscle co-activations seen after stroke. This hypothesis makes two predictions: the reticular formation should be naturally activated mainly for movements that compose the flexor synergy, and its outputs should coactivate muscles in a flexor-synergy-like pattern.

In this study performed in two healthy macaque monkeys, we set out to test whether neurons were preferentially activated for movements aligned to the flexor synergy. Importantly, we not only recorded from all major motor cortical regions, but also from the reticular formation. Surprisingly, we found that all brain regions studied – including the reticular formation – contained cells which could code independent contraction around elbow or shoulder joints. In addition, where cells were activated for co-contraction around these joints, the majority fired with movements orthogonal to the flexor synergy (e.g. shoulder abduction with elbow extension). Finally, juxta-threshold microstimulation of motor cortex or reticular formation never led to co-activation of muscles needed to produce shoulder abduction with elbow flexion; this was only seen when stimulating the spinal cord. Our results suggest post-stroke synergies do not arise because of inherent flexor biases in upregulated reticulospinal pathways. Instead, they point instead towards spinal circuits as the source of these biases. We propose that after stroke, contralateral corticospinal control is lost to varying degrees leaving no option but to route residual descending commands via these biased spinal pathways. This necessarily produces the flexor synergy.

## Results

### Behavioral Task

Monkeys performed a task requiring isometric contractions around the shoulder and elbow joints. The animal sat with the right forearm in a close-fitting cast attached to a six-axis force-torque transducer (Fig. 1A). Output from the transducer was used to calculate torques exerted by the shoulder and elbow joints. A cursor on a screen in front of the monkey was moved left-right by the shoulder abduction/adduction torque, and up-down by the elbow flexion/extension torque (white circle, Fig. 1B). After a period at rest, a single target appeared in one of 16 locations (squares, Fig. 1B); the monkey moved the cursor to the target and held to obtain a food reward. Figure 1CD show average shoulder and elbow torques from a single behavioral session, aligned to the start of the 0.4 s-long hold period (grey shading in Fig. 1CD). The different plots are arranged according to different target locations; for example, the top right plot reflects the target requiring a combination of shoulder abduction and elbow flexion. The torques were highly tuned by the nature of the task. For comparison, Fig. 1EF presents the average of rectified EMG from the shoulder abductor muscle posterior deltoid, and the elbow flexor biceps. As expected, these show very similar directional tuning to the shoulder and elbow torques (see polar plots in the middle of each panel, Fig. 1CDEF).

**Figure 1.**
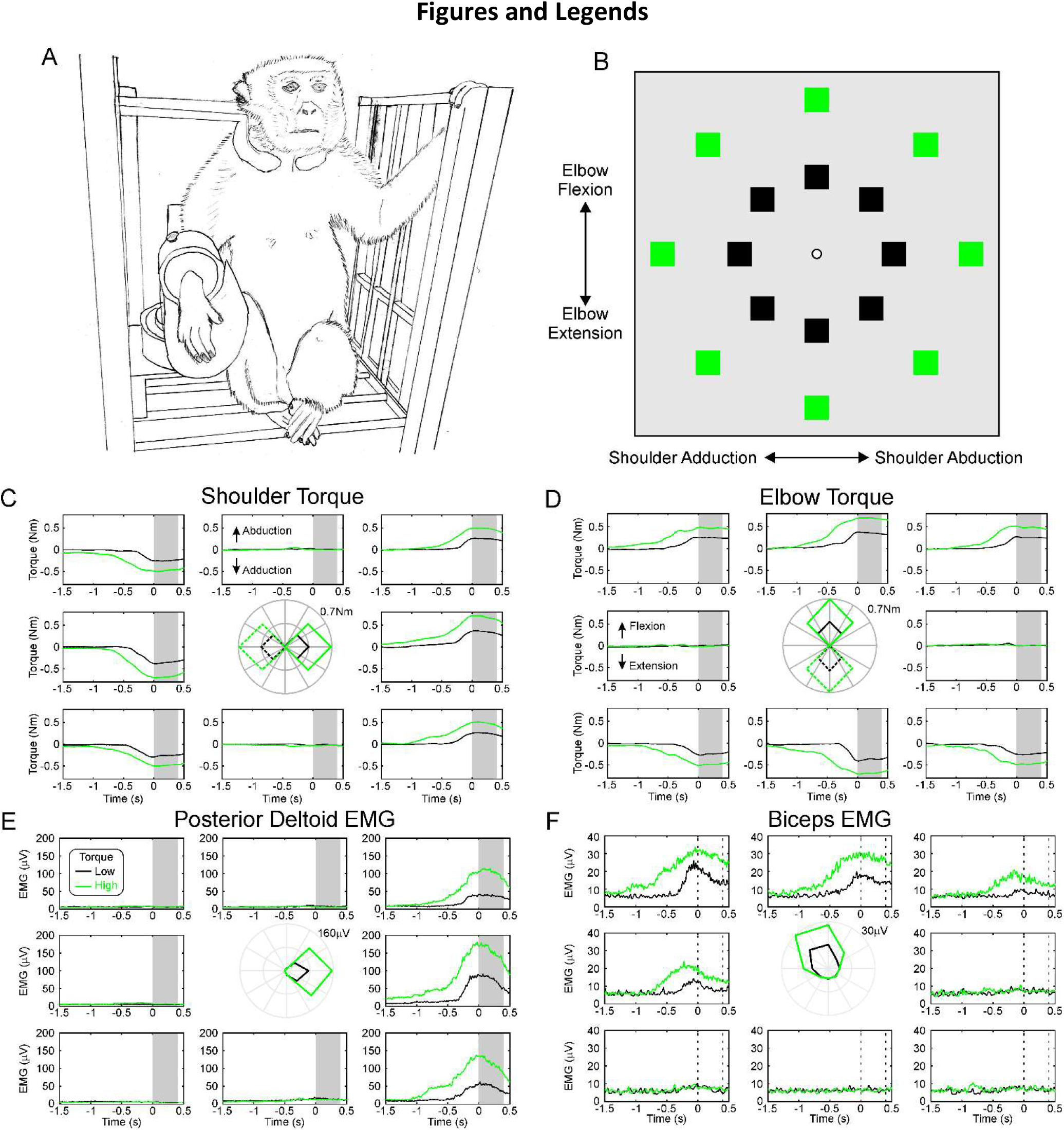
Behavioral Task. A, drawing of the experimental arrangement. The monkey’s arm was placed in a closely fitting cast linked to a six-axis force-torque transducer. B, screen arrangement viewed by the monkey. The central cursor (white) was moved left-right by shoulder abduction-adduction torques, and up-down by elbow flexion-extension torques. Squares show targets (only one appeared at a time). Green/black color marks targets close to/further away from the center (rest) position. C, shoulder, D, elbow torques. Each plot is positioned according to the location of targets on the screen. Traces are an average of all trials performed in one session in monkey C, aligned to the start of the hold period (shaded grey). Line colors indicate targets closer or further from the center, as in (B). Inner polar plot shows the torque as a function of target angle: solid lines indicate elbow flexion or shoulder abduction, dotted lines elbow extension or shoulder adduction. E, dependence on task of EMG from the posterior deltoid muscle, F, same for biceps muscle. Same arrangement of plots as (C,D).

### Single Unit Recordings

Peri-event time histograms (PETHs) were compiled of spikes relative to either the beginning of the task hold period, or to the onset of contraction. Figure 2A shows such PETHs for one neuron recorded in the reticular formation of monkey C, relative to the onset of contraction. PETHs are arranged similarly to the plots of Fig. 1. This neuron was strongly directionally tuned, preferring targets requiring shoulder adduction and elbow flexion; firing was slightly larger for the high *vs* low torque targets (black and green lines in Fig. 2A).

**Figure 2.**
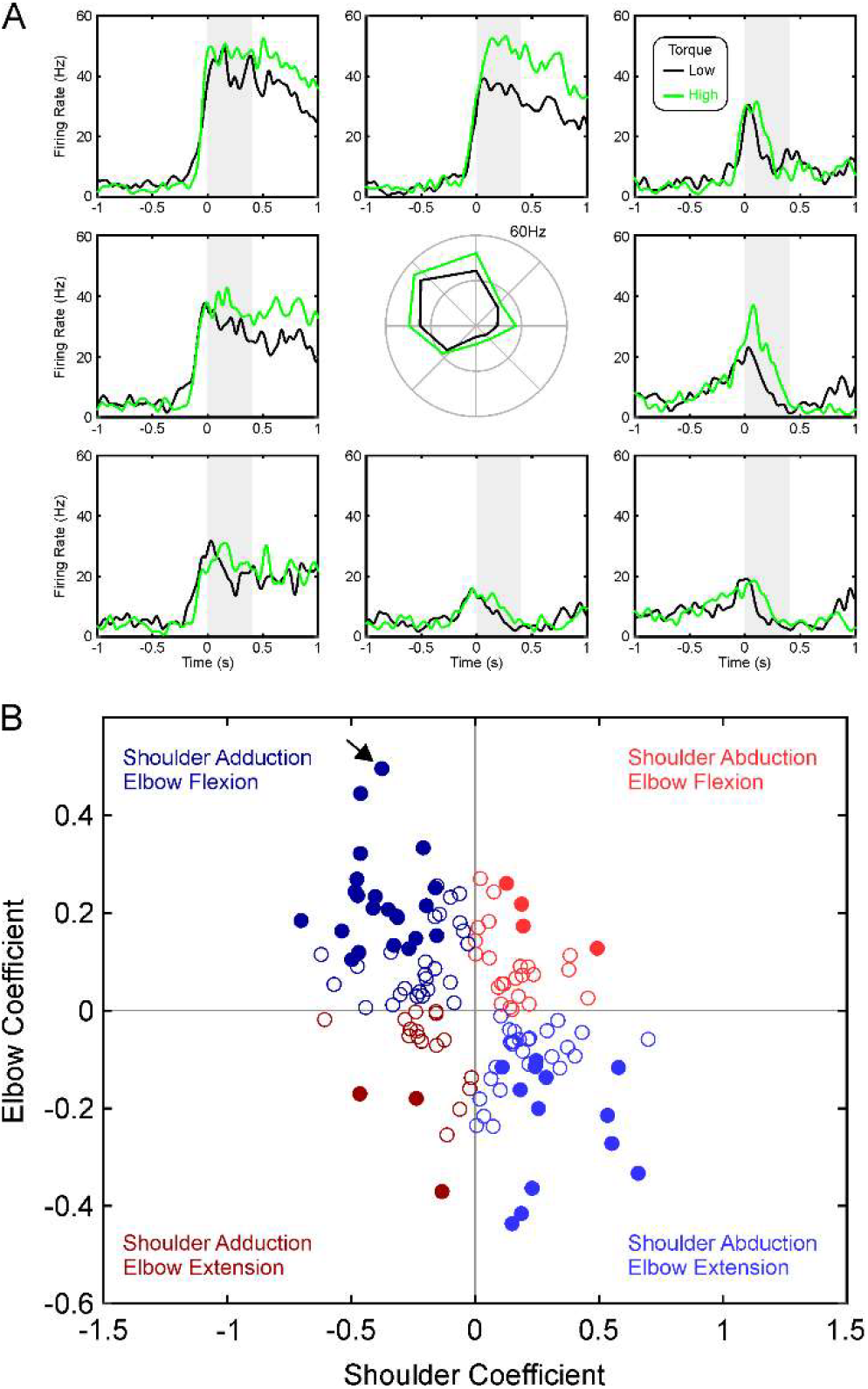
Example of Single Unit Activity. A, firing rate of a single unit recorded from the reticular formation in monkey C, using similar display conventions as Fig. 1. Alignment is to the onset of torque. B, scatter plot of regression coefficients for task-related cells recorded in monkey C from the reticular formation, for firing rates during the period shaded grey in A. Filled circles mark cells which were significantly related to both shoulder and elbow torques; open circles were only significantly related to torque from one joint. Colors indicate the four quadrants, corresponding to different combinations of elbow and shoulder torques. Red colors are *in synergy*, blue *out of synergy* as defined in the text. Arrow marks the cell illustrated in (A).

Mean firing rate was measured after the torque initiation (grey shading, Fig. 2A; values plotted using polar plot in Fig. 2A) and subject to regression analysis with torque as described in Methods, yielding regression coefficients. Figure 2B gives a scatter plot of these coefficients across all cells recorded in monkey C reticular formation. Firstly, we determined whether the coefficients relating firing to shoulder and elbow torque were significantly different from zero. This allowed classification of cells into those that were related to shoulder torque only, elbow only, both or neither. For the cells which were related to both shoulder and elbow torques (filled circles in Fig. 2B), further classification could be made based on what combination of torques they responded to (i.e. which quadrant they lay in on Fig. 2B). We designated cells related to shoulder abduction with elbow flexion, or shoulder adduction with elbow extension, as coding *in synergy* contractions (red colors in Fig. 2B). These cells had preferred combinations of torques which aligned to the pathological synergies seen after stroke. By contrast, cells which were related to shoulder abduction with elbow extension, or shoulder adduction with elbow extension, were designated *out of synergy* cells (blue colors in Fig. 2B) – because their preferred torque combinations were orthogonal to what is seen in stroke survivors.

### Neural Coding of Shoulder and Elbow Contractions

Figure 3 presents the number of cells which were placed into the different categories outlined above on the basis of regression analysis, for firing during the contraction phase of the task, where PETHs were aligned to onset of torque increase. Data from each animal are shown separately (left and right columns), and results are separated by the brain region (rows). The purple/green histograms show the numbers of cells coding for shoulder only, elbow only, both together or neither. The red/blue histograms show further details about the cells identified as coding for both elbow and shoulder torques (purple slices in the left pie charts), indicating whether they fired *in* or *out of synergy* as defined above.

**Figure 3.**
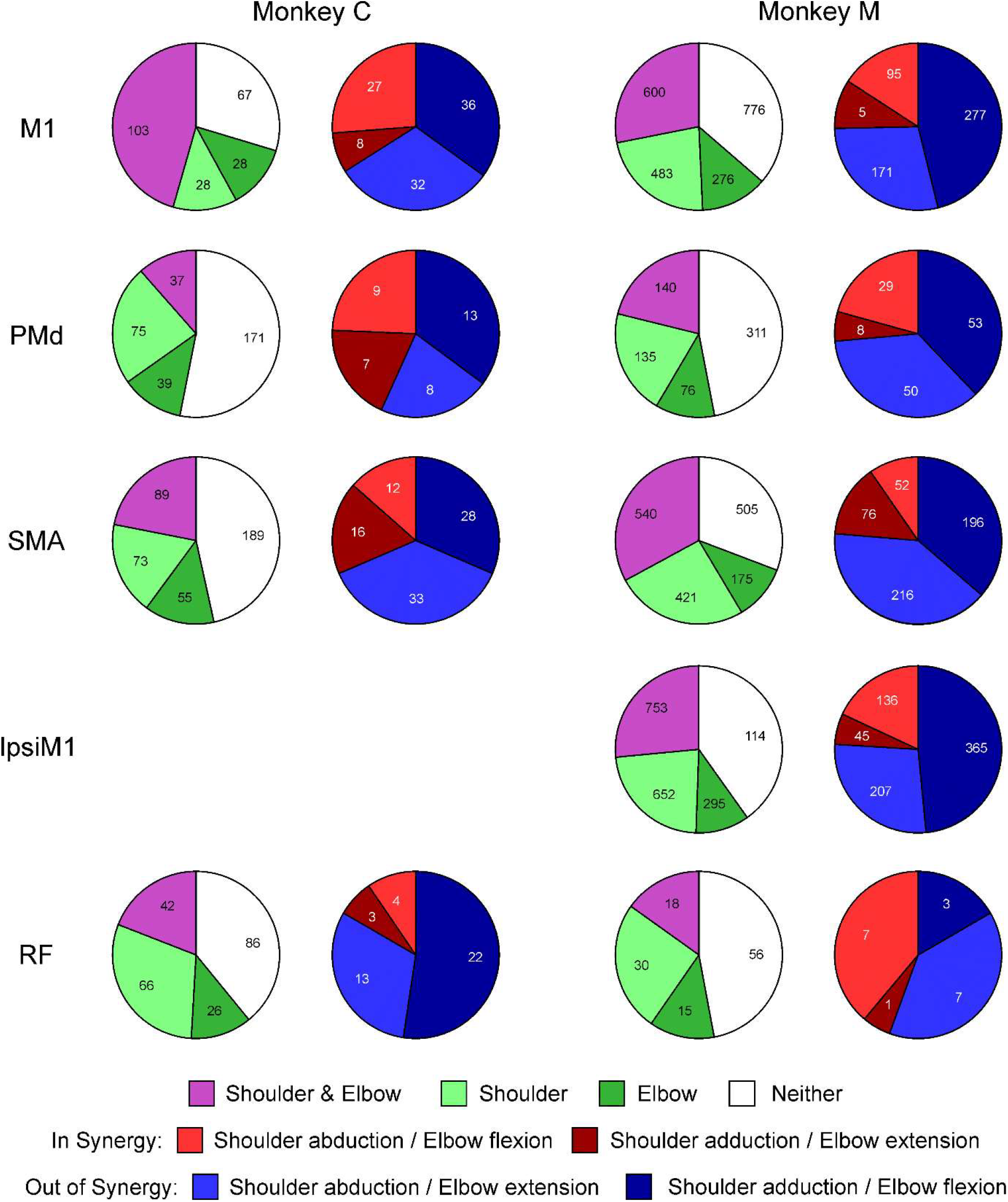
Distribution of Cells with Different Characteristics Related to Onset of Torque. Each row relates to a different region of the brain. Purple/green pie charts show the number of cells detected via regression analysis to be coding for torques around both shoulder and elbow joints (purple), only a single joint (different shades of green), or not coding for torque (white). Red/blue pie charts take cells included in the purple sector, and further classify them according to the type of co-contraction which they represent. Red colors indicate *in synergy* combinations; blue colors indicate *out of synergy* combinations. Results are shown separately for each monkey.

One hypothesis for the cause of post-stroke synergies is that they are produced when the loss of M1 circuits leads to control being taken over by another area – for example the ipsilateral M1 and/or the reticular formation. If that area were unable to produce independent control of the shoulder and elbow joints, but instead was constrained to produce synergistic co-contractions, this would lead to pathological synergies. Such synergies would not be seen in healthy individuals because M1 is able to counteract this tendency. Two aspects of the data in Fig. 3 argue against such a hypothesis.

Firstly, in all regions examined, there were many cells which coded only for elbow or shoulder torque (green sections in Fig. 3). Indeed, in all areas except M1 in monkey C, the cells coding for independent joint contraction outnumbered those coding for co-contraction (compare purple vs green sections). If any of these regions were to take over control following M1 damage, there is no reason to expect that they could not subserve fractionated joint control.

Secondly, when examining cells which do code for co-contractions, there were many cells which coded for out-of-synergy contractions – in all cases, these were in the majority. If synergies resulted from the release of a pre-existing bias in one of these regions, we would expect stroke survivors to show shoulder abduction coupled with elbow extension, or shoulder adduction coupled with elbow flexion – a pattern orthogonal to that actually seen. The bias in all areas was similar to that in M1. Comparing the proportions of *in synergy* to *out of synergy* cells in all areas relative to M1, only ipsiM1 and RF in monkey M were significantly different (chi-squared P=0.012 and P=0.0054 respectively; both significant after Benjamini-Hochberg correction for multiple comparisons). These differences were in opposite directions – ipsiM1 had fewer, and RF more *in synergy* cells, although the number of cells in the sample for RF was very low. However, importantly the majority of cells coding for co-contraction were aligned to *out of synergy* contractions in all areas.

Some previous work has suggested that the neural control of holding (maintaining fixed state) may engage different circuits from the control of movement (producing a change of state)^16-19^. Accordingly, it is possible that different results could be seen when analyzing holding versus movement components of a task. However, for the present study that does not seem to be the case. Figure 4 presents proportions of neurons from different areas in the same format as Fig. 3, but now for analysis of firing in the hold phase of the task. Very similar results are seen: all regions have cells coding for single joint contraction as well as for co-contraction, and of the latter cells are usually more numerous coding for *out of synergy* than *in synergy* co-contraction. When comparing the distribution of *in synergy* vs *out of synergy* cells for each monkey and area to that seen in M1, only ipsiM1 in monkey M was significantly different (chi-squared test P=0.0049, significant at P<0.05 after correction for multiple comparisons using Benjamini-Hochberg procedure) – and this difference was in the direction that ipsiM1 was even more biased than M1 to *out of synergy* movements. There was a strong correlation between the coefficients determined from torque initiation and those from the hold phase of the task (mean correlation r=0.69, range 0.42-0.9).

**Figure 4.**
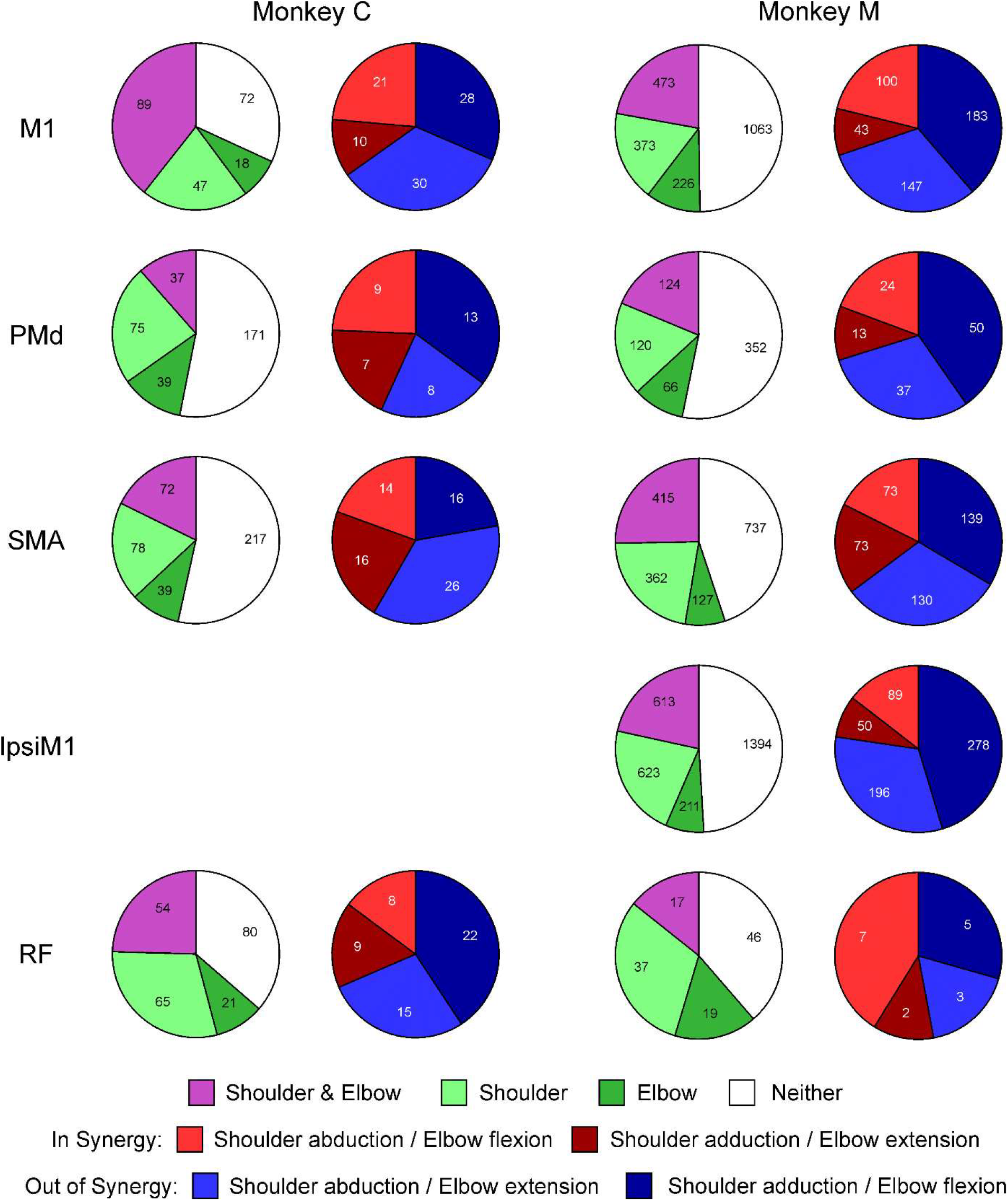
Distribution of Cells with Different Characteristics Related to Hold Period of Task. Similar display as Fig. 3, but for cells classified using firing measured during hold period.

The analyses of Figs 3&4 grouped cells into gross anatomical areas; we were interested to see whether similar results were seen with more nuanced sub-divisions. This is presented in Fig. 5; here results have been combined across animals; left and right columns present data for contraction and hold parts of the task respectively.

**Figure 5.**
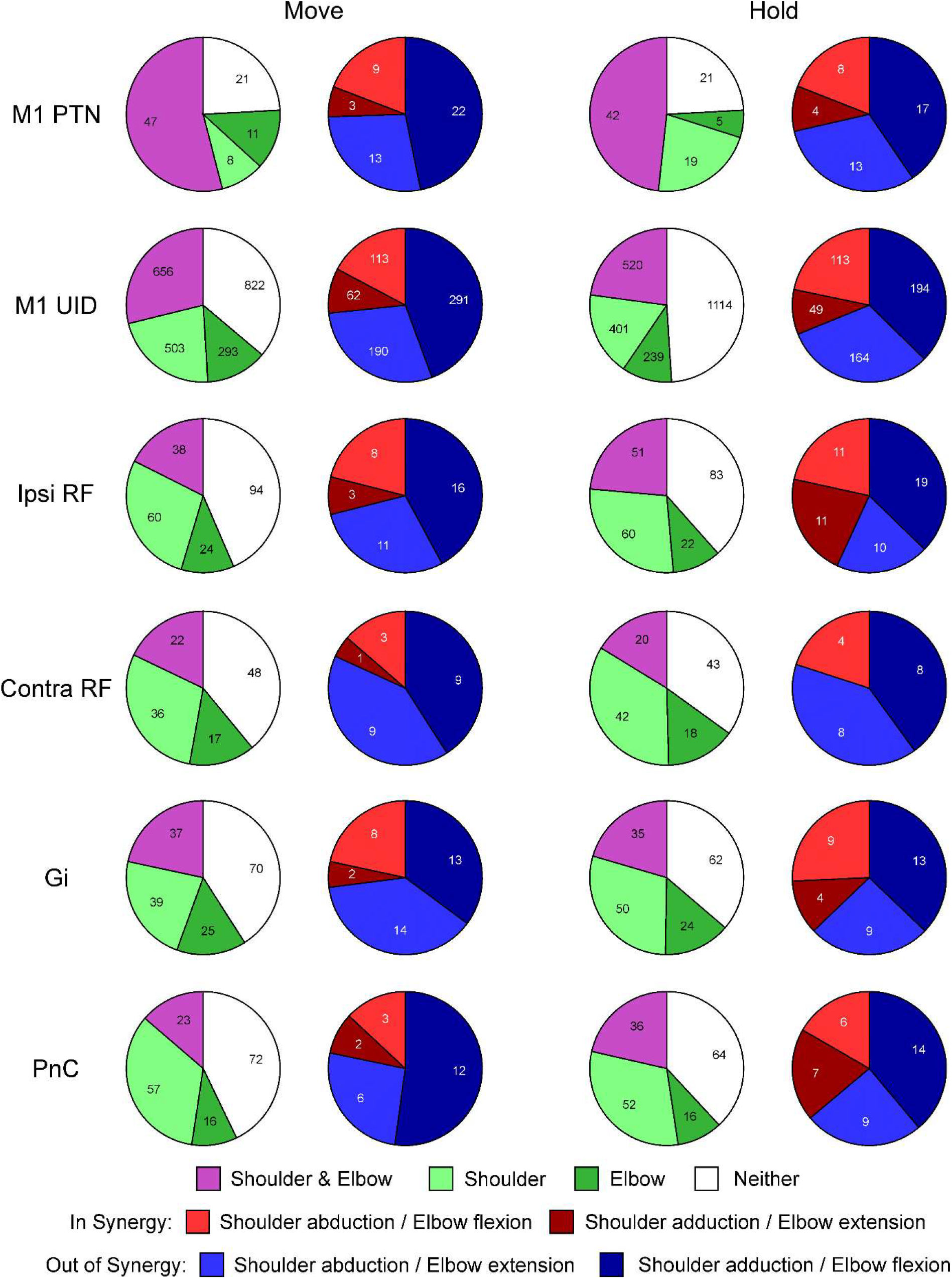
Distribution of Cells with Different Characteristics Separated by Cell Type. Results presented in Figs 4 and 5 for M1 and RF are separated by the class of cell. For M1, results are presented for identified pyramidal tract neurons (PTNs) or unidentified cells (UID). For RF, cells have been classified in two ways. IpsiRF/Contra RF separates the neurons according to which side of the brainstem they were recorded on (red vs black dots in Fig. 3). Gi/PnC separates cells according to their estimated rostro-caudal location, in nucleus gigantocellularis or the caudal part of the pontine reticular formation (see maps in Fig. 3). Results are combined across the two monkeys; data from contraction onset (Move) and Hold phases of the task are presented on the left and right side of the figure respectively.

In M1, we divided cells according to whether they were antidromically identified as PTNs which project to the corticospinal tract, or not so identified (unidentified cells, UID). Similar results were seen for both classes, although a higher proportion of PTNs were task related than UIDs (sum of purple and green segments).

The RF is a bilaterally organized structure. Reticulospinal cells receive bilateral input from the cortex^20^, and projections target both sides of the spinal cord^13, 21^. In this study our penetrations targeted RF both ipsilateral and contralateral to the limb performing the task (see Supplementary Figure 1). Figure 5 presents the results separately for each side; the findings are very similar. Finally, we separated cells depending on their estimated location within the brainstem, assigning them either to the pontine nucleus PnC or the medullary nucleus Gi. Once again, the results were very similar between the two nuclei: both contained cells coding for independent joint movement as well as for co-contraction, and more cells coding for co-contraction were activated *out of synergy* compared to *in synergy*.

### Muscle Responses to Microstimulation

Examination of cell discharge patterns during voluntary movement is a powerful approach, but conclusions are always based on correlation. This does not necessarily reflect the causal effect a cell has on muscles. A complementary approach is to measure muscle responses following weak electrical stimulation within a given area. This indicates the outputs sent from the stimulated site to spinal motoneurons, either via direct or indirect pathways. Here we analyzed responses evoked by microsimulation of M1, RF and also the intermediate zone of the spinal cord, focusing on whether regions were capable of supporting the pathological synergies seen post-stroke.

From all sites stimulated, at all tested intensities, we first selected only the instances where responses were produced in shoulder abductor or elbow flexor muscles. Responses were then classified according to whether they activated only one of these muscle groups, or both. The results are shown in Fig. 6 separately for M1, RF and spinal cord (SC) (rows). The left column includes all instances of recorded responses, regardless of stimulus intensity. Here, a concern could be that stronger stimuli spread and activated multiple outputs, making co-activation appear more common. The right column therefore includes only the smaller number of responses elicited at threshold, defined as the lowest intensity which elicited a response in any muscle.

**Figure 6.**
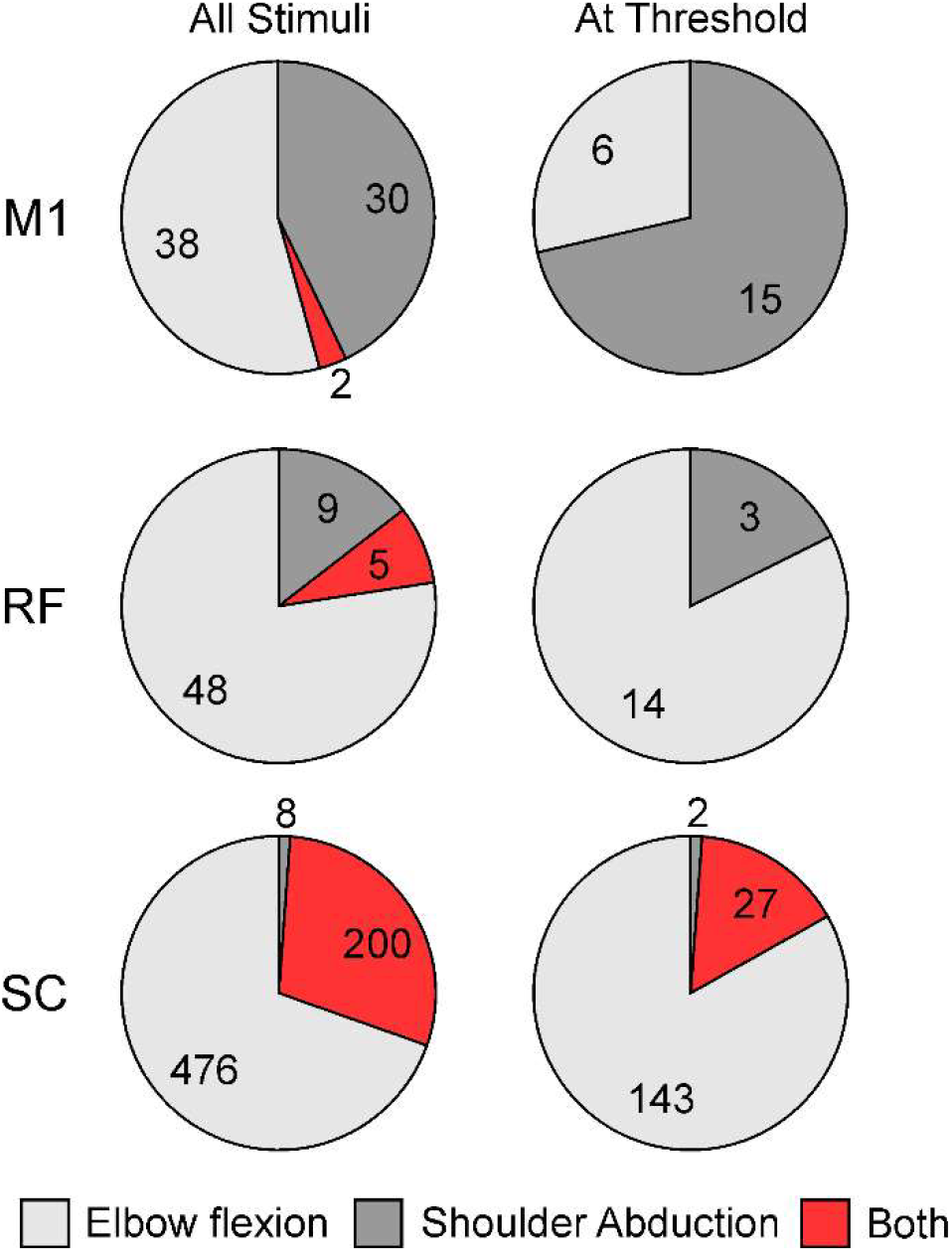
Effects Produced by Microstimulation. Each pie chart shows the distribution of effects produce by microstimulation in M1, RF or the spinal cord (SC). Effects are classified according to whether they activated elbow flexors only, shoulder abductors only, or both. Left column plots results for all available data, recorded at a range of stimulus intensities. Right column shows results only for sites stimulated at juxta-threshold stimulus intensity.

The overwhelming majority of sites within M1 and RF produced independent activation of either shoulder abductors or elbow flexors, but not both – at threshold, no sites were seen which produced co-activation of these muscle groups. By contrast, in the spinal cord at threshold only 2/172 sites produced independent shoulder abduction. The great majority of spinal sites (27/29, 93%) capable of activating shoulder abductors did so in consort with elbow flexion.

## Discussion

### Possible Sources of Post-Stroke Synergies

In this study, we show that cells within M1 can encode contractions around either the elbow or shoulder joint, or for co-contractions around both joints simultaneously. Furthermore, the cells which encode co-contractions have a bias: most cells code for contractions orthogonal to the synergies typically seen in a human stroke survivor. Considering these results from M1 in isolation leads to an obvious proposal for the source of post-stroke synergies: an M1 lesion removes drive for independent and *out of synergy* movements, leaving *in synergy* co-contractions as the only remaining option. However, critically here we recorded not just from M1, but from other cortical motor areas as well as the reticular formation. We show that all other brain centers tested had similar activity patterns to M1 (Fig. 3-5). Whichever one of these areas might take over function after M1 lesion, it should also be capable of providing drive for independent movements around single joints, and for across joint co-contractions the dominant drive should still be for *out of synergy* movements. Given these results, it is hard to propose that synergies arise because a pre-existing preference for *in synergy* co-contractions is unmasked in any surviving brain area. This is contrary to current ideas in the field, which suggest that after a severe lesion control is subsumed by ipsilateral motor areas and the reticular formation^10, 22-24^. These are proposed to be unable to encode fractionated independent contractions of elbow and shoulder joints^10^. Yet in our data, findings from ipsilateral M1 and the reticular formation were very similar to those from the contralateral M1, PMd and SMA.

Even if a brain region can accurately code for independent contractions across shoulder and elbow joints, these cannot be executed unless output projections are sufficiently well separated between different muscle groups. We investigated this by examining muscle responses to stimulation (Fig. 6). Juxta-threshold stimuli in both M1 and RF activated either elbow flexors or shoulder abductors, never the two groups together. M1 and RF not only have the ability to code for independent muscle contractions, but also to generate this independent activity via their output projections.

How then do post-stroke synergies arise? It is clear that something must change after the lesion. In health M1 corticospinal output probably does not provide the majority of drive to motoneurons^25^, but its input is still substantial. Part of this is via direct cortico-motoneuronal connections, as well as through oligosynaptic routes involving spinal cord interneurons^26-28^. Loss of M1 after a stroke therefore produces loss of motoneuron drive. Other pathways must then race to make good this deficit, sprouting terminals to strengthen their outputs. For the RF, electrophysiological recordings have shown that the majority of this additional input is disynaptic^15^. Anatomical studies also suggest that most increased corticospinal drive to the spinal cord after a lesion goes to lamina VI/VII, where interneurons are located; these provide a disynaptic route to influence motoneurons (from SMA bilaterally^29^ and from ipsilateral M1^30^). As Fig. 6 shows, in healthy animals spinal circuits almost always activated shoulder abductor muscles together with elbow flexors. A command attempting independent shoulder abduction from surviving ipsilateral cortex or the reticular formation would necessarily have to pass through these spinal interneurons, producing the undesired co-contractions characteristic of post-stroke synergies.

In many stroke survivors, synergies are only a transient phenomenon. With time, the ability to generate independent activity around shoulder and elbow joints can recover^1^. There are multiple possible ways in which this could occur. Monosynaptic connections to motoneurons, both from surviving cortical areas and from the RF, could increase, allowing fractionated control over muscles. Alternatively, spinal interneurons could refine their connections to target motor nuclei acting around only one joint. These options are not mutually exclusive. Importantly, some people are never able to go through this process, and remain stuck with unwanted synergies. These tend to be individuals with more severe lesions. We know that cortico-motoneuronal axons from M1 are exquisitely finely targeted, connecting to just a few motor nuclei^31,32^. It is possible that the survival of even a small number of these cortico-motoneuronal connections from the lesioned hemisphere could provide a ‘teaching’ signal, which allows spinal circuits to be gradually sculpted towards improved fractionation via activity-dependent processes. A severe stroke produces a near-complete lesion of these critical cells, and could therefore render it impossible to pass beyond *in synergy* movements.

### Cell Activity is Biased Towards *Out of Synergy* Movements

In this study, we provide evidence for a remarkable bias in cell firing patterns over regions within the cortex, as well as in the reticular formation. For neurons encoding contractions around both elbow and shoulder joints, the majority was related to shoulder abduction/elbow extension or shoulder adduction/elbow flexion (*out of synergy*)– the opposite of the obligate *in synergy* co-contractions seen post-stroke.

Somewhat different biases in cortical directional activity have previously been reported. In the classic study of Georgopoulos et al^33^ using a center-out reaching task in two dimensions, cell preferred directions were not uniformly distributed across the possible reach directions: reaches away from the body and to the right were over-represented. This would correspond to shoulder abduction with elbow extension (an *out of synergy* contraction); however, the opposite direction (towards body and left) was under-represented, which would require shoulder adduction and elbow flexion (also *out of synergy*). Such cell activity biases depend on the posture of the upper limb; with the shoulder abducted, reaches both towards and away from the body are over-represented^34,35^. This M1 preference is similar to that seen in muscle activity. It has been suggested based on limb biomechanics that towards/away movements require more power to produce a constant hand velocity than those to the left or right, leading to M1 distributing more resources to enable such movements^35^. By contrast, a study using three dimensional reaching found that preferred directions were approximately uniformly distributed^36^. None of these results are directly comparable to the present work, where we tested isometric contractions. Our task isolates motor control from any requirements imposed by limb kinematics. By focusing on the coordination of shoulder adduction/abduction and elbow flexion/extension, we tried to reveal control processes especially relevant to the development of obligate synergies post-stroke.

### Spinal Circuits are Biased towards *In Synergy* Movements

In contrast to our neural recording results from the cortex and RF, stimulation of the spinal cord showed a clear bias towards activating shoulder abductors with elbow flexors – an *in synergy* contraction. The spinal cord is known to play an important role in locomotion^37^, which muscles are activated in stereotyped synergies^38-40^. In quadrupedal macaques, both climbing and walking require shoulder abductors to be contracted with elbow flexors. The preference of spinal circuits for an *in synergy* movement may reflect a role best suited for producing locomotion. Descending control of a voluntary *in synergy* movement could be easily accomplished simply by activating the relevant spinal interneuron populations. By contrast, if the spinal cord cannot coordinate the relevant motor pools for an *out of synergy* movement, control would rely on more direct input from descending pathways. This could explain why more resources are devoted in cortical areas and the RF to control *out of synergy* vs *in synergy* movements. On this view, the development of descending pathways has been driven by the need to break away from the stereotyped co-contractions which subserve locomotion to a more flexible motor repertoire capable of supporting prehension. For any given movement, the relative contributions of primitive spinal circuits versus more selective descending pathways will depend on how well aligned the movement is to the capabilities of the spinal machinery alone.

To some extent, the widespread similarity between biases across the regions recorded from is unsurprising, as they are heavily interconnected. SMA and PMd both provide cortico-cortical input to M1^41-43^ and ipsiM1^44,45^. The RF receives highly convergent bilateral cortico-reticular input^20,46,47^. These interconnections should mean that the system is robust to damage; we would not expect, for example, that preferences of the RF would change extensively after damage to M1, as inputs to it from the SMA with the same biases would survive. These results, in addition to the novelty of suggesting that the post-stroke flexor synergy arises in spinal cord, also suggest that the cortex and brainstem evolved a control system to uncouple the arm from spinal circuits dedicated to locomotion, thus allowing prehension.

## Methods

Experiments were conducted on two female macaque monkeys (monkeys M, 7 years, 5.7 kg and C, 6 years, 6.1 kg). All procedures were approved by the Newcastle University Animal Welfare and Ethical Review Board, and were conducted under appropriate licenses from the UK Home Office.

### Behavioral Task

Animals were trained to perform a task requiring precise control of torque around the elbow and shoulder joints, which was measured using methods described previously^48^ adapted to be suitable for use in monkey. During training sessions, the monkey sat in a training cage, and was lightly restrained by a collar around the neck. A 3D scan of the right forearm was taken (EinScan Pro HD, SHINING 3D, Hangzhou, China), and used to construct a customized plastic cast by 3D printing. This was in two pieces which could be assembled around the forearm and then fitted into a circular clamp, providing a tight-fitting attachment which was without uncomfortable pressure points. The clamp was connected to a six axis transducer (model 45E15A4-I63-EF-200N, JR3 Inc, Woodland, CA, USA) which measured force and torque exerted by the animal (see Fig. 1A). The transducer and neck restraints were placed in the same locations on the training cage each day. Taking the center of the transducer as the origin of the coordinate system, we measured the 3D locations of the elbow and shoulder relative to this origin, and expressed these as vectors <u>*r*_*e*_</u> and <u>*r*_*s*_</u>. The force and torques measured in 3D by the transducer are the vectors <u>*F*</u> and <u>*M*</u> respectively. Analyzing the arm as a rigid body in equilibrium^49^, in which forces and torques are only applied at the shoulder and forearm attachment to the transducer, showed that the shoulder torque could be calculated as:

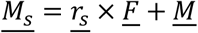

Where × denotes the vector cross product. The upper arm and forearm were then analyzed as two rigid bodies in equilibrium, in which the upper arm was acted on only by shoulder and elbow joints, and the forearm by the elbow and transducer attachment. This showed that the elbow torque could be calculated as:

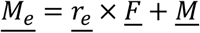

Voltage signals proportional to *<u>F</u>* and *<u>M</u>* were sampled by a PC with data acquisition card (National Instruments model USB6343, Austin, TX, USA), and the calculations described above performed in real time by a custom computer program. This displayed a cursor on screen. Movement of the cursor in the x axis was controlled by shoulder abduction-adduction torque. Movement in the y axis was controlled by elbow flexion-extension torque (see Fig. 1B).

Each trial of the task began with the cursor hidden. To proceed, the six force-torque signals from the transducer had to have a standard deviation (measured over a 1.3 s window for monkey C; 1.8 s for monkey M) below a threshold value (0.45 N for force, 0.05 Nm for torque); we found that this was a reliable indicator that the monkey was at rest. The force-torque signals were then zeroed; this compensated for slow drifts in the outputs which otherwise occurred over the course of a one-hour long training or recording session. The cursor then appeared, along with a target. Targets were spaced radially around two circles at 45° intervals, with the circle radius corresponding to a torque of 0.35 and 0.7 Nm. Targets were rectangular, with a half-width of 0.1 and 0.08 Nm in the shoulder abduction/adduction and elbow flexion/extension direction respectively (see Fig. 1B). The monkey was required to place the cursor into the target, and to hold for 0.4 s in monkey C, and 1 s in monkey M. The target and cursor then disappeared, and the animal was given a food reward. Placement of the cursor into target, and the successful completion of a trial, were indicated with auditory tones in addition to the on-screen information. The 16 different targets were presented in pseudorandom order in blocks of five.

The monkeys became highly skilled at performing this task. Monkeys C and M completed an average of 564 and 575 successful trials per recording session (range 236-880).

### Surgical Implants

After behavioral training was complete, the animals underwent an implant surgery. This implanted a headpiece, which permitted atraumatic head fixation and incorporated chambers to give access to brain regions with microelectrodes. The headpiece was an anulus generated by 3D printing in titanium, using a digital model derived from an MRI scan of the head; this ensured a good fit between the headpiece underside and the bone. The headpiece was attached to the skull using a system of expanding bolt assemblies as described previously^50^. Both the headpiece and bolt assemblies were coated in hydroxyapatite, to improve osteointegration. In the same surgery, fine stainless-steel wires insulated with Teflon (part number AS632, Cooner Wire Co, Chatsworth, CA, USA) were implanted in the following 12 muscles acting on the upper limb: extensor digitorum communis, extensor carpi radialis, flexor digitorum superficialis, flexor carpi radialis, triceps, brachioradialis, biceps, brachialis, posterior deltoid, anterior deltoid, supraspinatus, pectoralis major. These wires ran sub-cutaneously to connectors on the headpiece, allowing EMG recordings.

In a subsequent surgery, two fine electrodes (LF501, Microprobes Inc, Gaithersburg, MD, USA) were implanted in the pyramidal tract in the medulla, using a double angle stereotaxic approach^51^. Electrodes were advanced while stimulating (up to 500 μA isolated biphasic constant-current stimuli, 0.1 ms per phase); they were fixed in place at the location with the lowest threshold to produce an antidromic response in a recording made from the dural surface overlying the primary motor cortex (M1; thresholds at final location were between 15 and 145 μA).

All surgeries were conducted under general anesthesia. After induction with ketamine (7 mg/kg), medetomidine (3 μg/kg) and midazolam (0.3 mg/kg; all IM), anesthesia was continued with sevoflurane inhalation (2-2.25%) and alfentanil continual intravenous infusion (24 μg/kg/hr). The animal was intubated, and ventilated using positive pressure ventilation. Temperature was maintained with a supply of warm air, and a thermostatic heating blanket. Meloxicam (300 μg/kg) was given for added analgesia. Methylpregnisolone (loading dose 30mg/kg followed by 5.4 mg/kg/hr) was given to reduce cerebral oedema. Hartmann’s solution was infused to prevent dehydration (5 ml/kg/hr). Prophylactic antibiotic was administered (cefotaxamine, 20mg/kg every 4 hours). Monitoring included pulse oximetry, heart rate, core and peripheral temperature and non-invasive blood pressure. At the end of the surgery buprenorphine (20 μg/kg), amoxy-clavulanate (12.5mg/kg), dexamethasone (0.5 mg/kg), maropitant (1mg/kg) and paracetamol (23 mg/kg) were given.

### Neural Recordings

After recovery from the implant surgeries was complete, a craniotomy was opened under the recording chamber overlying M1 and the dorsal pre-motor cortex (PMd) in a further brief surgery, and recording sessions commenced. Electrodes were introduced through the dura, and moved to find single unit recordings. In monkey C, recordings initially used up to five glass-insulated platinum microelectrodes, positioned using an Eckhorn microdrive^52^; we searched particularly for antidromically-identified pyramidal tract neurons (PTNs). In later recordings, silicon probes with 32 contacts were used (Atlas Neuroengineering, Leuven, Belgium), followed by 1024 contact SiNAPS active neural probes^53, 54^. In monkey M, only 1024 channel SiNAPS electrodes were used in M1/PMd. For the multiple contact probes, recordings following pyramidal tract stimulation were made at the start and end of each recording session; this allowed us to identify antidromically identified PTNs^53^. Recordings were also made from SMA contralateral to the arm performing the task, from RF bilaterally, and (in monkey M only) from M1 ipsilateral to the task. All cortical recordings used SiNAPS probes, other than some sessions in M1 for monkey C as noted above. All RF recordings used 32 channel U probes (Plexon Inc, Dallas, Texas, USA).

### Histology and Cell Localization

At the end of neural recordings, the animals were used in an unrelated study under terminal anesthesia, and then killed by overdose of anesthetic and transcardial perfusion. The brain was removed and processed for histology. Parasagittal sections of the cortex surrounding the central sulcus and the brainstem were cut on a freezing microtome (50 μm thickness) and stained using cresyl violet. For the cortical sections, the boundary between M1 and PMd was determined as the point where large Betz cells ceased. This was used together with the coordinates of cortical penetrations to classify recorded neurons as lying within M1 or PMd. For the brainstem, a parasagittal section was digitized, and the reticular nucleus gigantocellularis (Gi) and caudal pontine reticular nucleus (PnC) were identified relative to landmarks which were marked on a traced drawing^55^. The drawing was aligned with the MRI of the brain taken at the start of the study; this allowed us to register the histological tracing to stereotaxic coordinates, and thereby to reconstruct electrode penetrations using manipulator readings taken each day. Brainstem recording sites were classified as lying on either the left or right side of the brain, and within the Gi or PnC nuclei.

Supplementary Figure 1 shows the reconstructed location of reticular formation cells which were significantly related to some aspect of the task. Localizing our recordings was made more straightforward because we identified the abducens nucleus in some penetrations, from its characteristic discharge during voluntary lateral eye movements and the small lateral deviations of the eye caused during microstimulation. Sites where abducens was identified are marked on the reconstruction in blue. Based on previously-described divisions of the primate reticular formation^55^, we marked a boundary line between the nucleus gigantocellularis (Gi) and the caudal pontine nucleus (PnC). This allowed cells to be classified according to their estimated nuclear location; we also noted the side of the brainstem on which neurons were recorded, allowing classification as ipsilateral or contralateral to the arm performing the task (red vs black dots, Suppl. Fig. 1).

### Data Analysis

#### Torque Analysis

For each trial, the shoulder and elbow torques were smoothed with a Gaussian kernel (50 ms width parameter), and binned into 10ms bins. The torque path (position of cursor on the screen) for each trial was then plotted from the trial start until the end of the hold period. Target distance was defined as the straight-line distance between the cursor position at the trial start and trial end. A threshold for movement onset was calculated for each trial by taking 5% or 10% of the target distance, for targets furthest or nearest the center position respectively. Movement onset was then defined as the last point where the distance from trial start was less than this threshold. Trials were only included in the analysis if this point was in the correct direction, defined as being within ±90° of the target direction.

To facilitate a comparison between torques and cell firing rates, a mean torque value was calculated for the movement period (400ms window starting at movement onset, as defined above) and the hold period (400ms window starting at start of the hold period). Baseline torque was subtracted from each of these values (mean torque across first 50ms of each trial, when the cursor was held on the central target). Mean values were calculated separately for shoulder torque and elbow torque.

#### Spike Discrimination

Neural recordings were first separated into the discharge times of clean single units. For single microelectrode recordings, this used a custom cluster-cutting program (Getspike, SN Baker). For 32 channel silicon probes and 1024 channel SiNAPS probes, this used KiloSort4^56^. Clusters identified by Kilosort were further processed by the program Bombcell^57^; only units classified as ‘good’ by both Kilosort and Bombcell were used in further analysis. For SiNAPS probe recordings, an additional more time-consuming analysis extracted the spike times of identified PTNs, using filters matched to the identified spike shape^53^. Kilosort clusters which matched these spikes were deleted to avoid duplication. Finally, brainstem recordings using 32 channel linear probes (U probes) were clustered using Mountainsort^58^; a custom program further processed these clusters to remove those likely to be artifacts, and to combine clusters likely to originate from the same cell.

#### Single Unit Analysis – Cell Selection

Spike times were converted into a continuous estimate of firing rate by convolution with a Gaussian kernel^59, 60^ (width 50 ms). Cells were occasionally lost during recording. To prevent these periods with no spikes from artifactually reducing estimates of task-related firing, we identified blocks of ten consecutive trials with no spikes recorded; these trials were removed from the analysis for that cell. Peri-event time histograms (PETHs) were then constructed by averaging across all trials for each condition. A bin width of 50 ms was used, and PETHs were smoothed by convolution with a Gaussian kernel (width 50 ms). PETHs were constructed separately aligned to torque onset and the start of the hold period, and all subsequent analyses were performed separately for these ‘move’ and ‘hold’ PETHs.

Cells were only included in the analysis if they met the following three criteria. Firstly, there had to be a minimum of five trials for each condition. Secondly, baseline firing had to be consistent across all conditions. This was assessed by averaging firing rate for each condition for a baseline period from −1.5s to −1.0s, relative to the trial marker (move onset or start of the hold period). Cells were excluded if any of these baseline firing values lay outside ±5 median absolute deviations from the median of baseline firing for that cell. Thirdly, cells were only included in the analysis if their firing was task related. This was determined using a Monte Carlo method. For each trial, the inter-spike intervals were shuffled and the PETH recomputed. This process was repeated to produce 100 PETHs with shuffled data. The latency of peak firing was calculated from the unshuffled PETH by finding the latency of peak firing for each condition and taking the median of these values. For each condition, firing rate at this peak latency was compared between the real PETH and the shuffled PETHs. The firing of a cell was considered task-related if the real peak firing was greater than at least 95 of 100 of the peak firing rates from shuffled PETHs, for at least half of the conditions. Note that since these analyses were performed separately for PETHs aligned to movement and the hold period, different populations of cells were included in the ‘move’ and ‘hold’ analyses.

#### Correlation of Torque and Firing Rate

Although the window of interest for torque was fixed, corresponding to either the first 400ms after movement onset, or the first 400ms of the hold period, cell firing rate presumably modulates prior to this to drive the contraction. Furthermore, it is likely that the latency between firing rate modulation and torque output varies for each cell. We therefore identified a window of firing rate for each cell that best correlated to torque and used this in our analysis.

For each cell, we calculated the mean firing rate for each 400ms window starting −200ms to 0ms relative to the movement onset, progressing in 10ms steps. We then performed a regression between each of these firing rates and mean shoulder/elbow torque during the initial movement period (0-400ms after movement onset). We selected the firing rate window with the highest regression coefficient to include in the subsequent analysis, representing the period of cell activity that best correlated with torque. Firing rate, shoulder torque and elbow torque values were all normalized by their respective standard deviations, prior to performing a Tobit regression^61^. This method was chosen here to accommodate the left-censoring of the firing rate data - rate values cannot be negative. Confidence intervals were calculated for the regression coefficients. This analysis was repeated for the hold period.

The coefficients from the Tobit regression were used to classify the relation of the cell activity to torque. A cell was defined as having a significant correlation between firing rate and shoulder torque if the confidence intervals of the regression coefficients did not cross zero. This analysis was repeated for elbow torque, thereby allowing classification of cells as ‘shoulder only’ (significant relationship between firing rate and shoulder torque, but not elbow torque), ‘elbow only’, ‘both’, or ‘neither’. Examination of the sign of the correlation coefficients for ‘both’ cells allowed a further classification according to the type of co-contraction which the cell responded to. *In synergy* co-contractions were those aligned to post-stroke synergies: elbow flexion with shoulder abduction, or elbow extension with shoulder adduction. *Out of synergy* co-contractions were those orthogonal to post-stroke synergies: elbow flexion with shoulder adduction, or elbow extension with shoulder abduction.

To compare coefficients between data aligned to torque onset and the hold phase of the task, for each monkey, area and joint a cell was included if the Tobit regression coefficient was significantly different from zero for either task phase. We then calculated the correlation between torque onset and hold coefficients.

### Analysis of Published Data on Responses to Microstimulation

To complement results from the new experiments described above, we also carried out some re-analysis of a previously published study^12^. Full experimental details are given in the publication. Briefly, EMG was recorded from many muscles acting around the shoulder, elbow, wrist and digits in one arm. Microelectrodes were advanced into M1, RF and the spinal cord intermediate zone (SC). Experiments in M1 and RF were carried out in the awake state, with access to the brain provided with recording chambers as described above. For SC, the recordings were made in a terminal experiment under propofol, an anesthetic which we have found leaves spinal excitability relatively intact. Single-pulse stimuli were given through the microelectrode, and responses in EMGs recorded. For each site, and for each stimulus intensity, the post-stimulus averaged rectified EMG was tested for statistical significance relative to the baseline, and the muscle was classified as responding or not responding. Where a response was seen, the intensity was progressively reduced until it was abolished. The threshold intensity was then defined as the lowest intensity which produced a response in at least one muscle. In the post-hoc analysis carried out here, sites were only included if they activated at least one muscle out of posterior deltoid, anterior deltoid, biceps, brachioradialis and brachialis. Sites were then classified according to whether they activated shoulder abductors alone, elbow flexors alone, or both (i.e. at least one muscle in each category).

## Acknowledgements

The authors wish to thank Terri Jackson and Andrew Atkinson for animal training, Norman Charlton for engineering support, Rocio Palacios O’Connor for veterinary assistance, and Michelle Waddle for surgical theatre nursing.

---

**Supplementary Figure 1.**
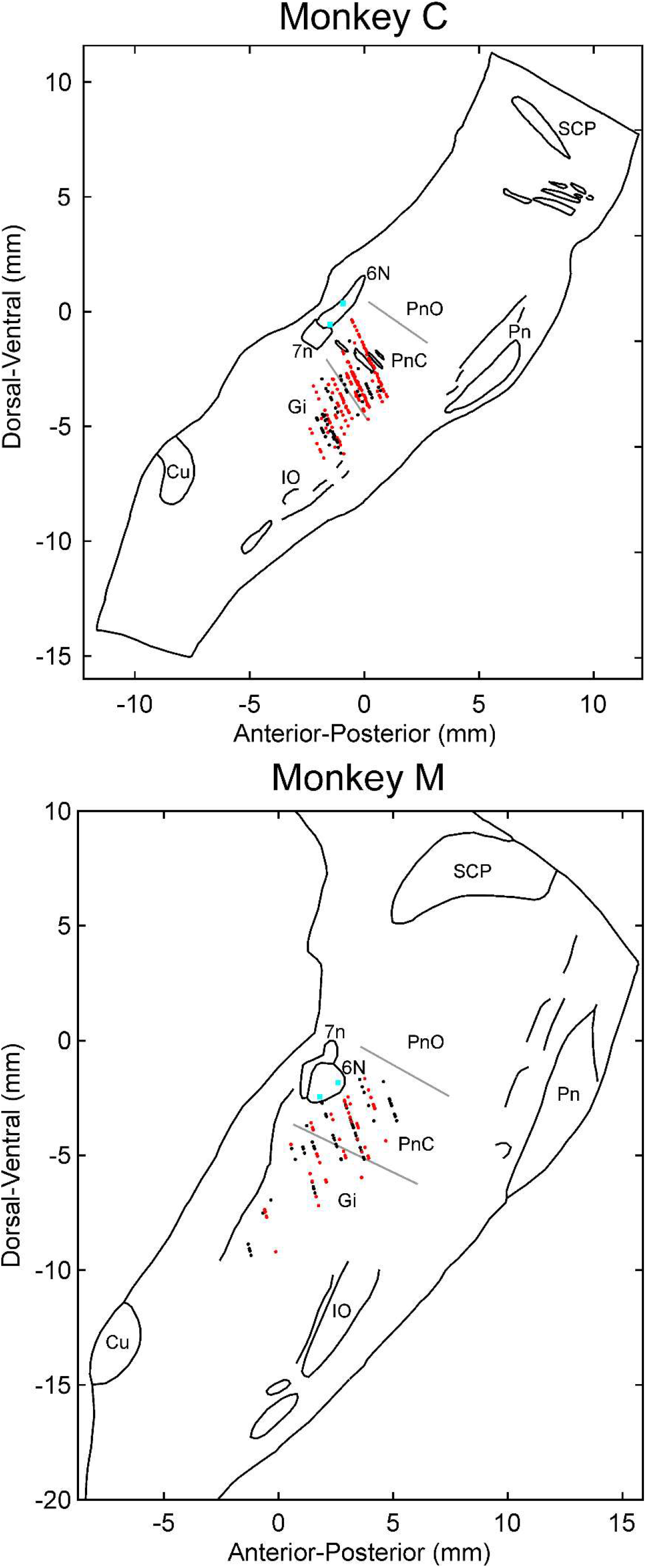
Reconstruction of Cells within the Reticular Formation. Estimated cell location is shown in stereotaxic coordinates (AP0 DV0 corresponds to the inter-aural line), superimposed on a tracing of a parasagittal histological section taken around 1.5 mm from the midline. Visible brainstem landmarks are indicated by labels. Only task-related cells are shown. Black dots mark cells on the left (contralateral), red cells on the right (ipsilateral) side of the brainstem. Blue squares plot locations where cells related to eye abduction movements were recorded, and/or eye abduction movements could be elicited by microstimulation. These sites within the abducens nucleus were used to align the map. IO, inferior olive. Cu, cuneate nucleus. 7n, facial nerve. 6N, abducens nucleus. SCP, superior cerebellar peduncle. Pn, pontine nucleus. Gi, nucleus gigantocellularis of the reticular formation. PnC, caudal part of the pontine reticular nucleus. PnO, oral part of the pontine reticular nucleus. Grey lines mark division between Gi, PnC and PnO used to classify cells.

